## Supplementary Materials for "Accounting for Isoform Expression in eQTL Mapping Substantially Increases Power"

#### 1 Preprocessing of GEUVADIS data

We obtained genotype and RNA-Seq data from the GEUVADIS [1] dataset (see Code and data availability in the main text). For both our simulated dataset and real data analysis, we used the genotypes of 87 Yoruban donors in the GEUVADIS data. We used kallisto [2] version 0.46.0 to quantify the number of reads mapped to each isoform using the RNA-seq reads for the Yoruban samples, and discarded isoforms for which fewer than 50% of the samples had at least 5 reads mapped to that isoform.

For our real-data analysis, the transcripts per million (TPM) quantifications provided by kallisto were then normalized using the "DESeq2::estimateSizeFactorsForMatrix" function from the DESeq2 [3] R package. For our simulations, the original quantifications were discarded and new phenotype values were simulated according to the model presented in the main text.

#### 2 Preprocessing of GTEx data

We largely preprocessed the GTEx genotype and expression data in the way described in the Supplementary Materials for the GTEx v8 analysis [4]. The main differences were that i) we used only European samples to limit potential population stratification; ii) we performed low-expression filtering on isoform expression levels rather than aggregated gene expression levels; iii) normalization was applied to isoform expression levels rather than aggregated gene expression levels.

We obtained a GCT file containing isoform expression levels for the thyroid, a covariates file for the thyroid, and a population file listing the European samples from the GTEx portal website (see Code and data availability in the main text). We additionally obtained a VCF file containing the genotypes (Code and data availability). All files were subsetted to include only samples from the thyroid where covariates were available and the donor had European ancestry.

Genotypes were processed according to the GTEx v8 steps outlined in Section 2.3.4 of their Supplementary Materials [4]. Most of these steps apart from minor allele frequency (MAF) filtering at 1% had already been performed by the GTEx consortium prior to obtaining the VCF. The only difference in our pipeline is that we filtered out variants that were not genotyped in 100% of the samples. We used PLINK [5] v1.90b6.6 to perform the MAF and genotype call rate filtering. The resulting VCF had 10,489,762 SNPs compared to 10,726,114 reported in GTEx v8 [4].

Phenotypes were converted from GCT to BED format and annotated with the "Homo_sapiens.GRCh38.88.chr.gtf" file from Ensembl [6] (Code and data availability). We used this file instead of the GENCODE file used in the GTEx v8 analysis because the GENCODE file does not include isoform names, only gene names, whereas the Ensembl file does have the isoform names. The phenotypes were normalized with TMM [7], filtered for low expression, and then inverse-normal transformed as described in the GTEx v8 paper's Supplementary Material Section 3.4.2 [4]. Unlike in the GTEx v8 analysis, the phenotypes here were isoform expression levels, rather than gene expression levels.

### 3 Software details

We wrote our own implementations of the Wilks-Bartlett test [8], the Cauchy aggregation test [9, 10], and the minimum p-value method. For Fisher's method [11], we used the version implemented in the scipy [12] python library. We used QTLtools [13] version 1.3.1. We wrote our own implementation of the FastQTL "Beta approximation" permutation scheme [14] and used this permutation scheme for all methods except QTLtools, which has its own implementation of this scheme built-in. For q-value false discovery rate (FDR) control [15], we used version 2.20.0 of the "qvalue" R package.

Any gene with a q-value of less than 0.1 (for simulations and GEUVADIS analysis) or 0.05 (for GTEx analysis) was considered an eGene identified by the method being evaluated. For GEUVADIS data and simulations, all methods were run with cis-windows of 50kb on each side of the gene's transcription start site. For GTEx, all methods were run with cis-windows of 1Mb on each side of the gene's transcription start site, to match the GTEx v8 analysis [4]. All methods were set to not output results or perform permutations on SNPs that had a nominal gene-level association p-value of greater than 0.5. In principle, all SNPs should be used to compute the p-values used in the permutation test described in the FastQTL paper [14]. However, in practice, this filtering step barely impacts results, since these SNPs have such low association signal, and results in a large speedup.

The Cauchy aggregation test [9] can in principle incorporate weighted p-values. Since the sample size is consistent across isoforms, we implemented the unweighted version of this statistic. It is possible that weights could be employed for some other purpose, e.g. if prior information is available, but this is outside the scope of the present study.

### 4 Covariance simulation details

Most of the parameters of the simulation are described in the main text. Here, we describe how $\mathbf{\Phi}_j$ – the covariance matrix for the effects of a cis-SNP on isoform expression levels – and $\mathbf{V}$ and $\mathbf{W}$ – the covariance matrices used for simulation of the matrix normal noncis effects – are simulated. For all three, we start by generating a matrix of the appropriate dimension where the diagonal entries are 1 and the off-diagonals are drawn from $Unif(\rho_{min}, \rho_{max})$, where $\rho_{min}$ and $\rho_{max}$ were set to 0.09 and 0.49 respectively for $\mathbf{\Phi}_j$ and $\mathbf{V}$, and were set to 0.99 each for $\mathbf{W}$.

This is not guaranteed to generate a positive semi-definite matrix, however. For $\mathbf{\Phi}_j$ we simply re-sample the matrix until it is positive semi-definite. For $\mathbf{W}$ this procedure is not tractable because the sample size is relatively large. Therefore, for $\mathbf{V}$ and $\mathbf{W}$, we sample a matrix $\mathbf{A}$ according to the procedure outlined above, then compute $\mathbf{A}^T\mathbf{A}$, which is guaranteed to be positive semi-definite. We then divided all entries in the resulting matrix by the maximum value in the matrix. We found that this generally resulted in diagonal values close to 1 and off-diagonal absolute values generally between 0.4 and 1 for $\mathbf{V}$ and generally between 0 and 0.2 for $\mathbf{W}$.

### 5 Functional enrichment analysis details

We performed functional enrichment analysis using torus [16] and the SNP annotations provided in the GTEx v8 dataset [4] (available via the GTEx portal; see Code and data availability), mirroring the analysis performed in the GTEx v8 paper. We chose QTLtools-sum as a representative gene-level method because it is the most similar approach to FastQTL [14], which is used in the GTEx v8 analysis. We chose the F-test as a representative isoform-aware approach because it had the highest empirical power among methods that never had an inflated false positive rate in any simulation setting. We ran a "nominal" pass for each of these methods on the GTEx v8 data, in which all SNP-gene associations passing the standard $5 * 10^{-8}$ Bonferroni-corrected p-value threshold were called as eQTLs and input to torus; we only included SNP-gene pairs for genes that were identified as eGenes by each method.

Torus requires t-statistics for each SNP-gene association as part of its input. However, QTLtools-sum does not provide association statistics as part of its output and the F-test returns F statistics instead of t-statistics. Instead, we generated t-statistics from the p-values via

$$t = 1 - g(p/2, dof)$$

where $t$ is the t-statistic we generate, $g$ is the t-distribution CDF, $p$ is the p-value, and $dof$ is the degrees of freedom, which we set to the sample size minus two. The sign of the statistic is not recoverable, but we found that flipping or randomizing the sign had no impact on the results.
